# The Regional Landscape of the Human Colon Culturome in Health and Cystic Fibrosis

**DOI:** 10.1101/2025.09.29.679198

**Authors:** Sarvesh V. Surve, Rebecca A. Valls, Kaitlyn E. Barrack, Lorraine L. Gwilt, Timothy B. Gardner, George A. O’Toole

## Abstract

Cystic fibrosis (CF) alters gut physiology, yet its impact on microbial communities across colonic regions (ascending, transverse, descending colon) and microhabitats (lumen, mucosa) remains incompletely understood. Here, we applied culturomics to characterize gut microbiota in 32 individuals (22 nonCF, 10 CF). Persons with CF (pwCF) exhibited significantly higher viable bacterial loads than nonCF individuals, particularly in mucosal samples. Anaerobes predominated overall, with relative enrichment of aerobes in the mucosa of pwCF. Alpha diversity was reduced in mucosal samples and aerobic cultures for pwCF, whereas beta diversity was influenced by all the tested variables except the colonic region. Phylum-level analyses revealed enrichment of Proteobacteria and depletion of Actinobacteria, Bacteroidota, and Firmicutes in samples from pwCF, consistent with stool analysis. Random forest models identified selected oral-associated microbes as key predictive taxa and accurately classified polyp status with very high accuracy. Whole-genome sequencing of *Bacteroides fragilis* (n=21) and *Escherichia coli* (n=15) isolates, representing a subset of 109 gut bacterial genomes sequenced from this cohort, revealed minimal genomic variation across colonic regions and sample types, indicating intra-individual strain stability. The understandings from this pilot culturome study may help in developing targeted microbial therapeutic approaches to address the gut dysbiosis of CF.

**Importance:** This pilot study represents the first culturome analysis of the cystic fibrosis colon. Our preliminary findings demonstrate that CF-associated gut dysbiosis is spatially specific, with mucosal bacterial communities showing pronounced alterations while luminal communities remain unchanged. This spatial specificity suggests the mucosal microenvironment as a potential therapeutic target and indicates that interventions focused solely on luminal bacteria may be insufficient. The promising predictive accuracy of culturome-based machine learning models in this small cohort suggests these viable bacterial signatures could serve as biomarkers for CF management pending larger validation studies. Additionally, our initial observations of complex CFTR modulator effects on gut microbial communities provide insights for future studies optimizing combination therapies.

## Introduction

Cystic fibrosis (CF) is a genetic disorder caused by mutations in the cystic fibrosis transmembrane conductance regulator (CFTR) gene, affecting approximately 70,000 individuals worldwide (1, 2). While CF is primarily recognized for its devastating respiratory manifestations, gastrointestinal complications also affect many persons with CF (pwCF) (2, 3). The gastrointestinal tract of pwCF is characterized by viscous mucus production, impaired epithelial ion transport, chronic inflammation, and altered digestive enzyme function, creating a unique microenvironment that profoundly influences microbial community composition and function (4–9). Recent advances in microbiome research have revealed that gut dysbiosis in pwCF is not merely a consequence of disease progression but represents a fundamental feature that emerges early in life with lasting implications (3, 10–13). Infants and children with CF exhibit distinct stool microbiota compared to nonCF controls, with alterations including reduced genera associated with immune programming that remain largely unchanging within individuals over time (10–13). More critically, a progressive fecal dysbiosis for infants with CF is specifically characterized by decreased Bacteroidetes and increased Proteobacteria and is associated with early linear growth failure (10, 11, 14). These early-life microbial alterations also establish a dysbiotic foundation that appears to persist throughout the lifespan of pwCF, suggesting that adult gut microbiome patterns may reflect the consequences of early microbial programming rather than simply acute responses to disease progression or treatment.

The CF microbiome extends beyond a single body site, with emerging evidence supporting complex interactions between gut, respiratory, and oral microbial communities (10, 14–16). Crosstalk between the gut and lung, termed as gut-lung axis, has been shown to play an important role for pwCF (16). Intestinal *Bacteroides* secreted propionate has been demonstrated to modulate systemic levels of inflammatory cytokines (17). The validation of the gut-lung axis suggests that alterations in one microbial niche can have systemic effects, potentially explaining aspects of the multiorgan manifestations characteristic of CF.

Most CF gut microbiome studies have focused on fecal samples, which primarily reflect luminal rather than mucosal-associated communities (18). The gastrointestinal tract exhibits significant spatial heterogeneity in terms of anatomy, physiology, and microbial composition. Moreover, the distinction between luminal and mucosal microbial communities is particularly relevant in CF, where altered mucosal properties and impaired epithelial function may selectively affect bacteria that interact directly with the intestinal mucosa. We initially attempted to study the mucosal and luminal microbiome of colonoscopy patients by 16S rRNA gene sequencing, but were unsuccessful due to the apparently low level of microbes in our samples. We next turned to culturomics. The culturome, defined as the collection of microorganisms that can be cultured under laboratory conditions, represents a complementary approach to non-culture based approaches; the culturome focuses specifically on viable, metabolically active bacteria (19). This methodology enables the isolation and characterization of living bacteria (especially if they are low-abundance), facilitates downstream functional studies, and provides insights into clinically relevant organisms that may serve as therapeutic targets or biomarkers.

In this study, we aimed to explore the regional landscape of the human colon culturome in pwCF and a comparator nonCF cohort. This work provides a pilot dataset for understanding how CF shapes the gut microbiota and highlights microbial features with potential clinical relevance.

## Materials and Methods

### Patient Recruitment and Procedures

Individuals undergoing colonoscopy at Dartmouth Hitchcock Medical Center (DHMC) were recruited for this study. Samples were collected from both pwCF and nonCF individuals without any exclusion criteria. Demographic and clinical metadata were recorded for each participant (Table S1, supplementary data). Colonoscopy aspirates and mucosal brushes were collected from the ascending, transverse, and descending colon of 34 individuals (22 nonCF, 10 CF, and 2 heterozygotes).

### Colonoscopy Preparation and Sample Collection

Participants underwent a standardized polyethylene glycol (PEG 3350; Miralax)-based preparation to ensure adequate bowel cleansing. Seven days before colonoscopy, participants discontinued iron supplements (multivitamins were withheld only if they contained iron) and were advised to consult their prescribing physician regarding anticoagulant use. Aspirin and pain medications were continued. Two days prior, participants began a low-fat, low-fiber diet, avoiding nuts, seeds, raw fruits/vegetables, and whole grains. The day before colonoscopy, participants adhered to a strict clear-liquid diet (including water, broth, clear juices, tea/coffee without dairy, clear sodas, and non-red/blue/purple gelatin or popsicles). The evening before the colonoscopy, participants ingested four 5 mg bisacodyl tablets. Participants consumed 32 oz of the solution (238 g of PEG 3350 dissolved in 64 oz of a clear sports drink (excluding red, purple, and blue formulations) in 8 oz aliquots every 15–20 minutes, followed by the remaining 32 oz consumed within 4-6 hours before colonoscopy. Bowel preparation was considered adequate when the effluent was clear or light yellow. Colonoscopy was performed under conscious sedation. During the procedure, mucosal brushes and luminal aspirates were collected under sterile conditions for culturomic and sequencing analyses. The samples were transported to the lab on ice and processed immediately, and aliquots were stored at -80 °C.

### Sample Processing and Culture Conditions

All samples were collected and processed as outlined in Figure 1A. Brush samples were suspended in 3 mL phosphate-buffered saline (PBS) and vortexed vigorously, while aspirates were processed directly. Six samples per subject (aspirates and brushes from ascending, transverse, and descending colon) were plated in duplicates on sheep blood agar (BA) plates for either colony-forming unit per ml (CFU/ml) enumeration or 16S rRNA gene amplicon sequencing. Remaining material was stored in aliquots with or without 30% glycerol at -80 °C. For CFU determination, samples were serially diluted in PBS and plated on BA plates. One set of each BA plates was incubated aerobically and the other anaerobically (GasPak, BD Ref No. 260678). Aerobic colonies were enumerated after 24 h, and anaerobic colonies after 48 h at 37°C. Statistical comparisons were done using the Wilcoxon test and mixed-effects linear models in R. For sequencing, 100 μL of the sample was spread on BA plates and incubated under the same conditions outlined above. After incubation, plates were scraped, resuspended in 1 mL PBS, and pelleted. DNA was extracted using the Zymo Quick-DNA Fecal/Soil Microbe Miniprep Kit (Cat. # D6010) following the manufacturer’s instructions.

**Figure 1.**
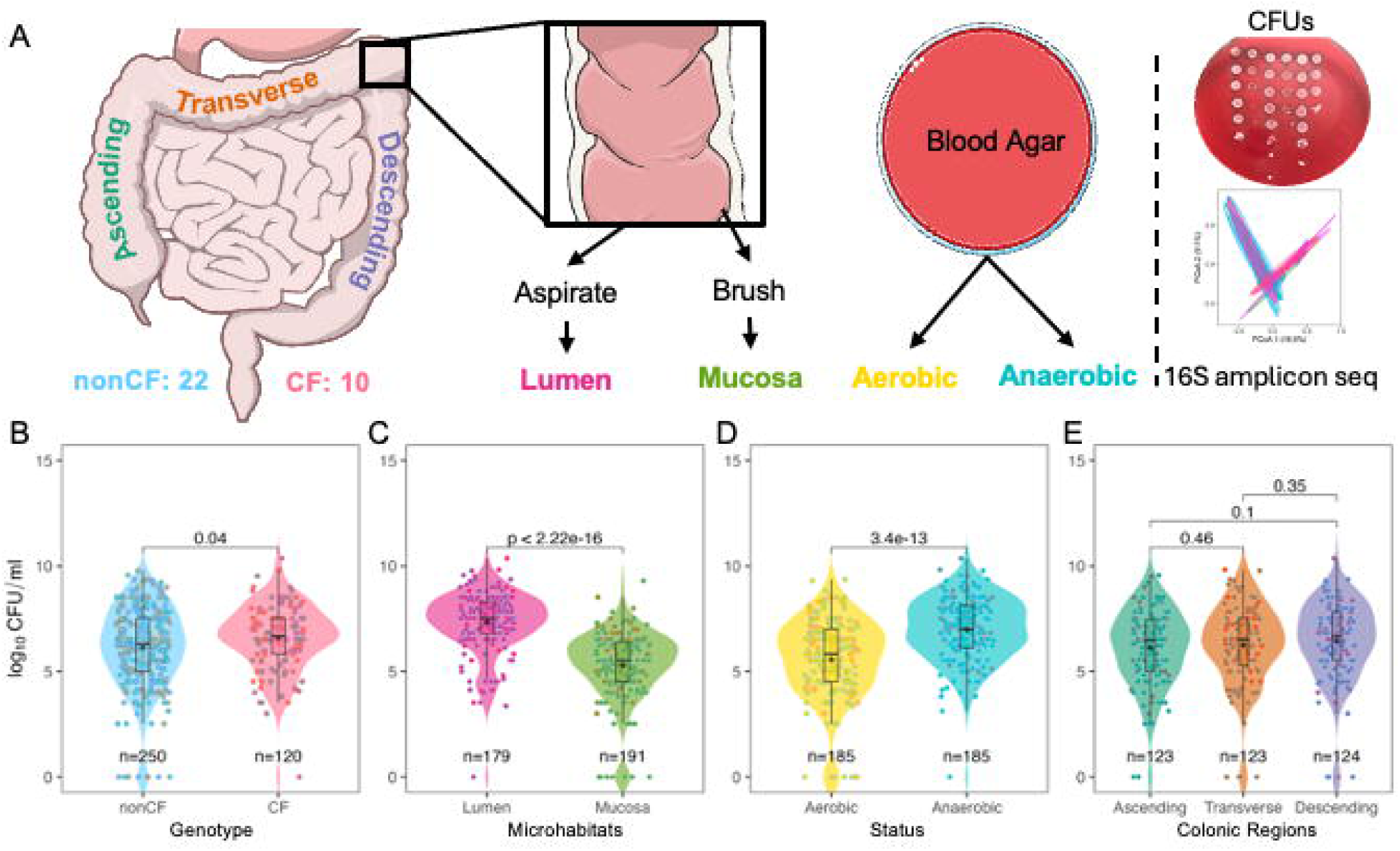
Overview and viable growth of colonoscopy samples. (A) Schematic of the study design. Luminal and mucosal samples were collected from the ascending, transverse, and descending colon of CF and non-CF patients. Samples were plated on blood agar for quantification of viable counts (log_10_ CFUs) and subjected to amplicon sequencing. (B–E) Viable counts (log_10_ CFUs) on blood agar comparing: (B) patient genotype (CF vs. non-CF), (C) sample microhabitat (luminal vs. mucosal), (D) oxygen status of growth conditions (aerobic vs. anaerobic), and (E) colonic region (ascending, transverse, descending). In all plots, black diamonds indicate group means. Colored dots are annotated by colonic location (color key: panel A) in panels B–D and by genotype in panel E. Statistical comparisons were made using the Wilcoxon test for each category.

### Amplicon Sequencing

The bacterial community composition was profiled by 16S rRNA gene amplicon sequencing of the V3–V4 hypervariable region. Library preparation and sequencing were performed at SeqCoast Genomics (Portsmouth, NH). Briefly, DNA extracted from cultured samples was subjected to PCR amplification using universal primers 341F (5′-CCTACGGGNGGCWGCAG- 3′) and 806R (5′- GGACTACNVGGGTWTCTAAT-3′) with Illumina overhang adapters.

Amplification was performed, followed by SPRI bead cleanup and subjected to indexing using Q5 High-Fidelity 2X Master Mix (NEB) and IDT for Illumina DNA/RNA UD Indexes. Indexed products were purified, pooled, and diluted samples were sequenced on the Illumina NextSeq 2000 platform, generating paired-end 2 × 300 bp reads. Raw sequence reads have been deposited in the NCBI Sequence Read Archive under BioProject accession number PRJNA1299459.

### Amplicon Data Processing and Bioinformatics

Raw paired-end reads were demultiplexed and processed using the DADA2 pipeline (v1.26) implemented in R (v4.2.1) (20). Reads were quality filtered with default parameters, including truncation based on Phred quality scores, removal of ambiguous bases, and filtering of reads with an expected error >2 for forward and >5 for reverse. Forward and reverse reads were denoised, dereplicated, and merged to reconstruct full-length amplicons. Chimeric sequences were identified and removed using the consensus method within DADA2. Amplicon sequence variants (ASVs) with a total abundance ≤10 reads across all samples were discarded to reduce spurious low-abundance variants. ASVs were taxonomically classified against the SILVA rRNA database (release 138.1) using the naïve Bayesian classifier implemented in DADA2. The resulting ASV tables were imported into phyloseq (v1.42) for downstream diversity and statistical analyses (21). Alpha-diversity (Shannon, Simpson, and observed ASVs) and beta-diversity (Bray–Curtis dissimilarity) metrics were calculated using phyloseq. Ordinations were performed by principal coordinates analysis (PCoA). Statistical significance of community differences by clinical variables was assessed using PERMANOVA (adonis2 in vegan R package, v2.6-4). Alpha diversity measures were compared using the Wilcoxon test and mixed-effects linear models in R.

To identify microbial features predictive of various clinical variables, we implemented random forest models using the *randomForest* R package (v4.7). Models were trained on genus-level centered log-ratio normalized count tables, with different clinical variables as the primary classification outcome. Feature importance was assessed using the mean decrease in accuracy metrics. For polyp predictions, data were randomly partitioned into training (75%) and test (25%) sets, and model performance was evaluated on the withheld test set. Features consistently ranked as highly informative across bootstrap replicates were considered candidate microbial biomarkers.

### Strain Collection and Genome Sequencing

The colonoscopy samples were streaked on different selective media: BA+ 100 μg/ml Gentamycin, *Bacteroides* bile esculin agar (BBE), and MacConkey agar (McC), and incubated both aerobically and anaerobically at 37 °C. The individual colonies were selected and streaked at least three times to obtain pure cultures. A subset of isolates from colonoscopy samples was sequenced for whole genomes. Sample libraries were prepared using the Illumina DNA Prep kit and IDT 10bp UDI indices, and sequenced on an Illumina Novaseq, producing 2x151 bp reads (UNH Hubbard Center for Genome Studies, Durham, NH) and Illumina NextSeq 2000, producing 2x151 bp reads (SeqCenter, Pittsburgh, PA, and SeqCoast Genomics, Portsmouth, NH) (22). Pangenome was assembled using anvi’o (v 8) (23).

## Results

To understand the differences in microbial populations across the colon, colonoscopy samples were collected from the ascending, transverse, and descending colon of 34 individuals: 22 nonCF, 10 pwCF, and 2 heterozygotes (Table S1, supplementary data) and the samples analyzed via a culturomics pipeline (Fig. 1A). The median age of the individuals was 51.5 years (range: 23-68 years). The differences in age between the CF and nonCF cohorts were nonsignificant, as determined by the Wilcoxon test. Two heterozygotes (carrying one CF and one nonCF allele) were excluded from culturomics analyses but retained for strain isolation. The BMI values were binned according to the NIH BMI calculator (24). The summary of subject-specific characters is given in Table 1. Among the 10 CF patients, 6 were on modulators and 5 had CFRD. All the pwCF on modulators were on the elexacaftor/tezacaftor/ivacaftor (ETI) combination. From each colonic region (ascending, transcending, and descending), two types of samples were obtained: mucosal (via brush) and luminal (via aspirate). Samples were plated on sheep blood agar (BA) for either colony-forming unit (CFU/ml) enumeration (Table S2, supplementary data) or amplicon sequencing (Fig. 1A).

**Table 1:** Summary of colonoscopy host-associated factors (summary excludes the heterozygotes). SD: standard deviation, PPI: use of proton pump inhibitors, NA: Not available. Details for each category are given in Table S1, supplementary data.

|  | **CF** | **NonCF** | **Total** |
| --- | --- | --- | --- |
| **Individuals, n (%)** | 10 (31.3%) | 22 (68.7%) | 32 (100%) |
| **# CFU, Amplicon** | 120, 119 | 250, 226 | 370, 345 |
| **Age in years, median ± SD** | 46.5 ± 16.1 | 53 ± 9.9 | 52.5 ± 12.5<br>(CF vs nonCF, p=0.19) |
| **Sex** | Female: 4; Male: 6 | Female: 14; Male: 8 | Female: 18; Male: 14 |
| **BMI** | <24: 4; 24-30: 2;<br>>30: 4 | <24: 8; 24-30: 8;<br>>30: 6 | <24: 7; 24-30: 15; >30: 10 |
| **Probiotics** | Yes: 3; No: 7 | Yes: 3; No: 19 | Yes: 6; No: 26 |
| **Antibiotics** | Yes: 6; No: 4 | Yes: 1; No: 21 | Yes: 7; No: 25 |
| **PPI** | Yes: 9; No: 1 | Yes: 6; No: 16 | Yes: 15; No: 17 |
| **Smoking** | Yes: 0; No: 10;<br>Former: 0 | Yes: 4; No: 15;<br>Former: 3 | Yes: 4; No: 25;<br>Former: 3 |
| **Alcohol** | Yes: 6; No: 4;<br>Former: 0; NA: 0 | Yes: 13; No: 6;<br>Former: 1; NA: 2 | Yes: 19; No: 10;<br>Former: 1; NA: 2 |
| **GI symptoms** | Yes: 8; No: 2 | Yes: 9; No: 13 | Yes: 17; No: 15 |
| **Pathology** | Yes: 6; No: 4 | Yes: 10; No: 12 | Yes: 16; No: 16 |
| **Polyps** | Yes: 3; No: 7 | Yes: 9; No: 13 | Yes: 12; No: 20 |
| **CFRD** | Yes: 5; No: 5 |  | Yes: 5; No: 5 |
| **Modulators** | Yes: 6; No: 4 |  | Yes: 6; No: 4 |

### Higher viable bacteria load in the CF colon mucosa

Overall, pwCF exhibited a significantly higher viable bacterial load than those without CF (median 6.7 vs. 6.3 log10 CFU/ml; Fig. 1B). Among the individuals with CF, those on modulators had a higher bacterial load (median 6.8 vs. 6.5 log10 CFU/ml; Fig. S1A). Luminal samples contained ∼100-fold more viable bacteria than mucosal samples (median 7.5 vs. 5.5 log10 CFU/ml; Fig. 1C). Across all samples, viable anaerobes outnumbered aerobes by more than 15-fold (7.0 vs. 5.8 log10 CFU/ml; Fig. 1D). Viable bacterial loads were comparable across colonic regions (ascending = 6.5, transverse = 6.5, descending = 6.7 log10 CFU/ml), although the transverse colon showed a non-significant trend toward higher counts (Fig. 1E).

When stratified by sample type and oxygen conditions, mucosal but not luminal samples showed distinct differences between CF and nonCF. CF mucosal samples contained significantly higher viable counts under both aerobic and anaerobic growth conditions (Fig. 2A). In contrast, luminal samples did not differ between CF and nonCF individuals, regardless of colonic location or oxygen availability (Fig. 2A, Fig. S1B). Within mucosal samples, aerobic viable counts in the descending colon and anaerobic counts in the transverse and descending colon of CF individuals trended higher, although these differences did not quite reach statistical significance (Fig. 2B).

**Figure 2.**
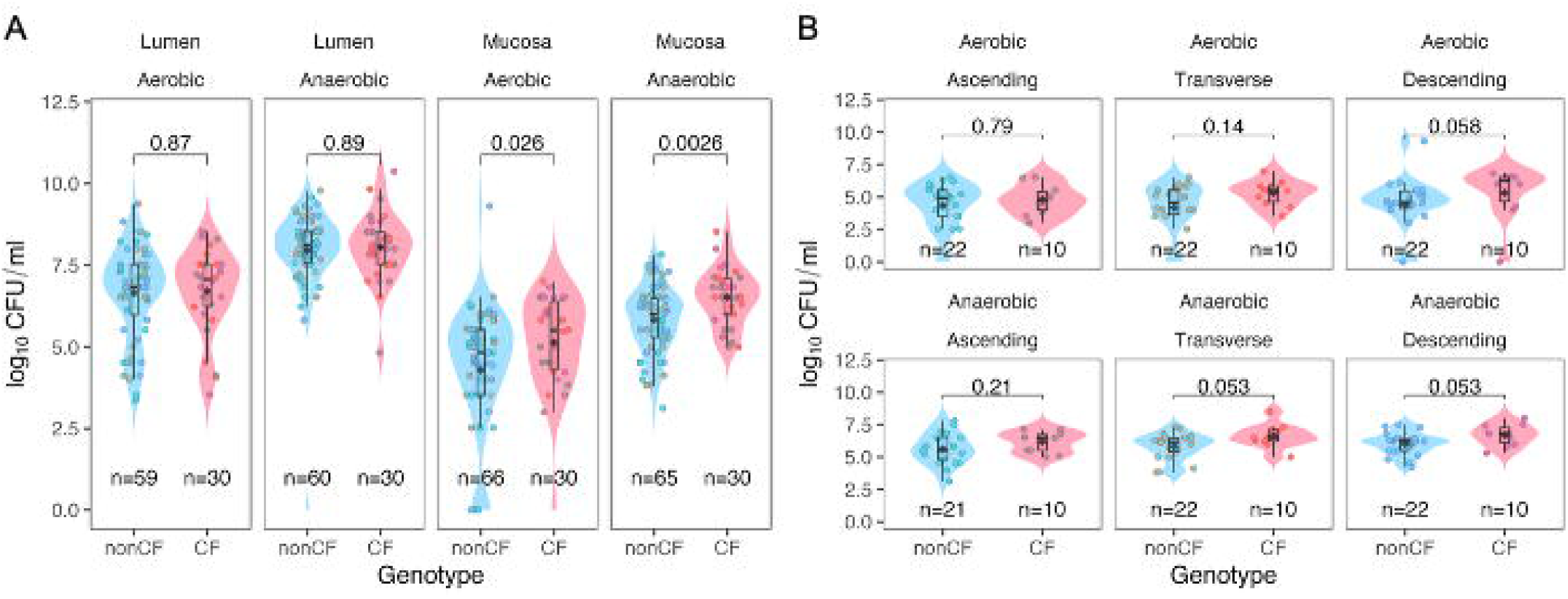
CF individuals have higher CFUs in mucosal-associated samples. (A) Viable counts (log CFUs) comparing CF and non-CF genotypes, faceted by microhabitat (luminal vs. mucosal) and oxygen status (aerobic vs. anaerobic). (B) Viable counts (log CFUs) for mucosal samples only, stratified by colonic region (ascending, transverse, descending) and oxygen status (top: aerobic; bottom: anaerobic). In all plots, black diamonds indicate group means. Colored dots are annotated by colonic location. Statistical comparisons were made using the Wilcoxon test for each category.

Among other tested variables, only the individuals with diagnosed pathology, such as the presence of adenomas and polyps, hyperplasia, and colitis, exhibited significantly higher viable bacterial counts (Fig. S1C). To account for repeated measures, we also performed a mixed-effects linear model with patient as a random variable, and most of the comparisons agree except for genotype (Table S3, supplementary data).

### Host and environmental factors shape culturome diversity

To investigate bacterial community composition of the culturome, we sequenced the 16S rRNA gene amplicon library of each sample and compared diversity across sampling location, culture condition, and patient variables. Alpha diversity was measured using the Shannon Diversity Index (SDI), Simpson Index, Chao1, and observed diversity (Table S4, supplementary data).

Comparisons across all the measures were similar; SDI was visualized as a representative ananlysis. SDI was significantly lower in pwCF compared to nonCF individuals across all samples (median 5.5 vs. 5.7 SDI; Fig. 3A), in mucosal compared to luminal samples (median 5.6 vs. 5.7 SDI; Fig. 3B), in individuals with antibiotic use (median 5.5 vs. 5.7 SDI; Fig. S2A), and in individuals with a BMI <25 compared to BMI ≥ 30 (median 5.6 vs. 6.0 and BMI 25-29.9: 5.5 SDI; Fig. S2B). All but one individual in the antibiotic use category belonged to the CF group as well, hence the differences we see might be due to the genotype. In contrast, anaerobically cultured samples exhibited significantly higher SDI than those grown under aerobic conditions (median 6.0 vs. 5.4 SDI; Fig. 3C). Higher SDI was also associated with alcohol use (median 5.7 vs. 5.5 SDI; Fig. S2C) and modulator therapy in pwCF (median 5.8 vs. 5.3 SDI; Fig. S2D). No significant differences in alpha diversity were observed across colonic regions, sex, probiotic use, PPI use, smoking, CFRD status, GI symptoms, or diagnosed pathology (Fig. 3D, Fig. S2E- K). The linear model on log-normalized SDI with the patient as a random variable showed only oxygen status and type of sample as significant variables (Table S5, supplementary data).

**Figure 3.**
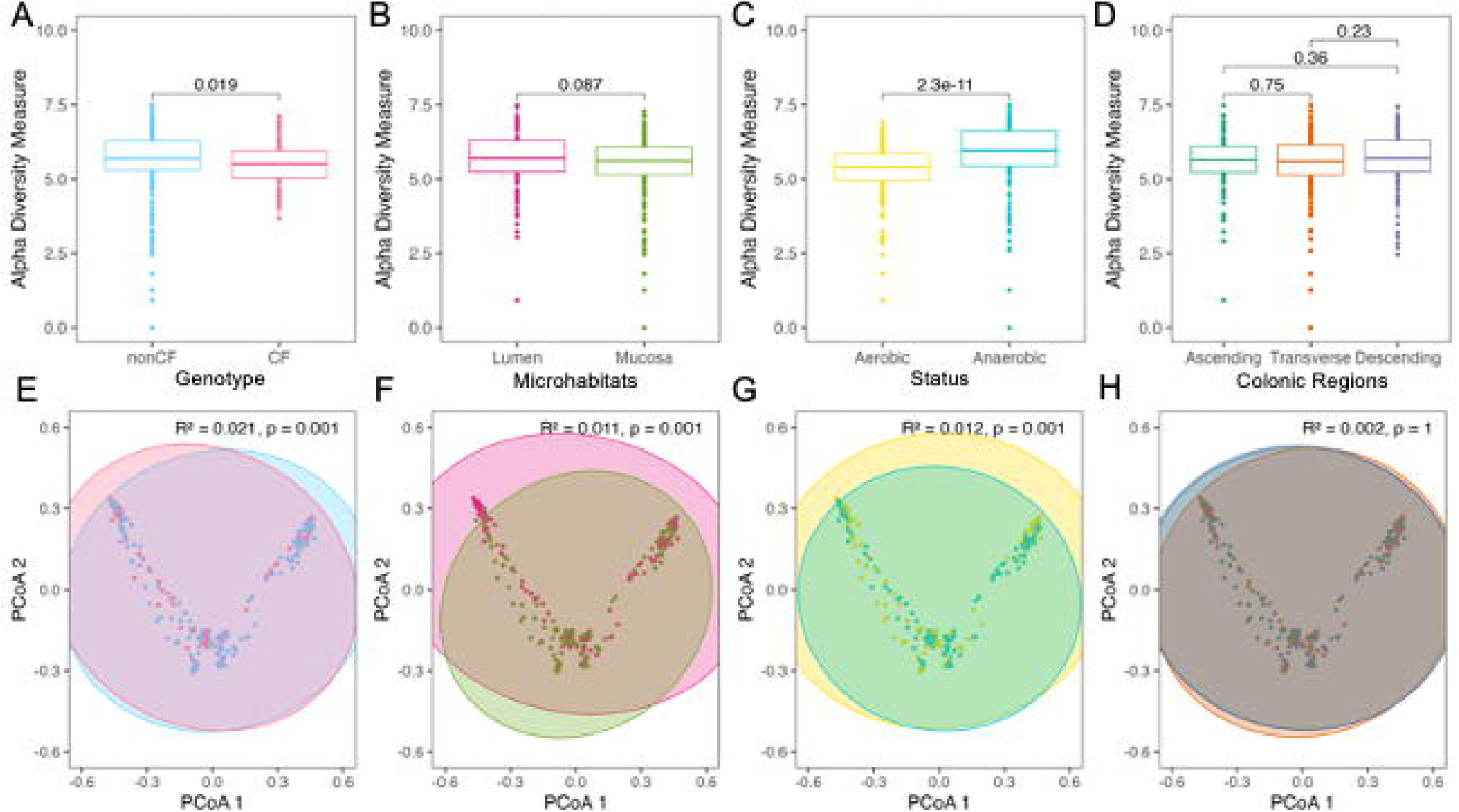
Anaerobic samples exhibit higher bacterial diversity. (A–D) Alpha diversity of all samples, calculated using the Shannon Diversity Index (SDI), and (E–H) Beta diversity, based on Bray-Curtis dissimilarity and visualized using Principal Coordinates Analysis (PCoA), comparing: (A, E) patient genotype (CF vs. non-CF), (B, F) microenvironment (luminal vs. mucosal), (C, G) oxygen status of growth conditions (aerobic vs. anaerobic), and (D, H) colonic region (ascending, transverse, descending). Statistical comparisons were performed using the Wilcoxon test for alpha diversity (A–D) and PERMANOVA for beta diversity (E–H).

Beta diversity assessed using Bray-Curtis or Morisita-Horn distances and tested with PERMANOVA, revealed similar values by both indexes and significant differences across nearly all variables except colonic location. By Bray-Curtis Index, the data only explain 31.9% of the total variance (R^2^) (Fig. S3A). Comparative plots are shown in Fig. S3B-L and Fig. 3E-H. Among all covariates, alcohol use (R^2^: 4.9%), BMI (R^2^: 4.7%), smoking (R^2^: 3.3%), modulator therapy (R^2^: 3.1%), antibiotic use (R^2^: 2.9%), genotype (R^2^: 2.1%), and probiotic use (R^2^: 1.9%), explained the greatest variation in community composition; however, together these factors accounted for only 22.9% of the total variance, indicating substantial individual variation not captured by measured variables.

### Phylum-specific alterations in CF culturomes

We next examined the taxonomic composition of all samples at the phylum level. Consistent with the diversity analyses, CF samples displayed reduced community diversity and appeared to be enriched in Proteobacteria compared to nonCF samples (median 48.4 vs 57.5% relative abundance; Fig. 4A). Community profiles stratified by other variables, including colonic regions and microhabitats as well as aerobes versus anaerobes, are shown in Figure S4.

**Figure 4.**
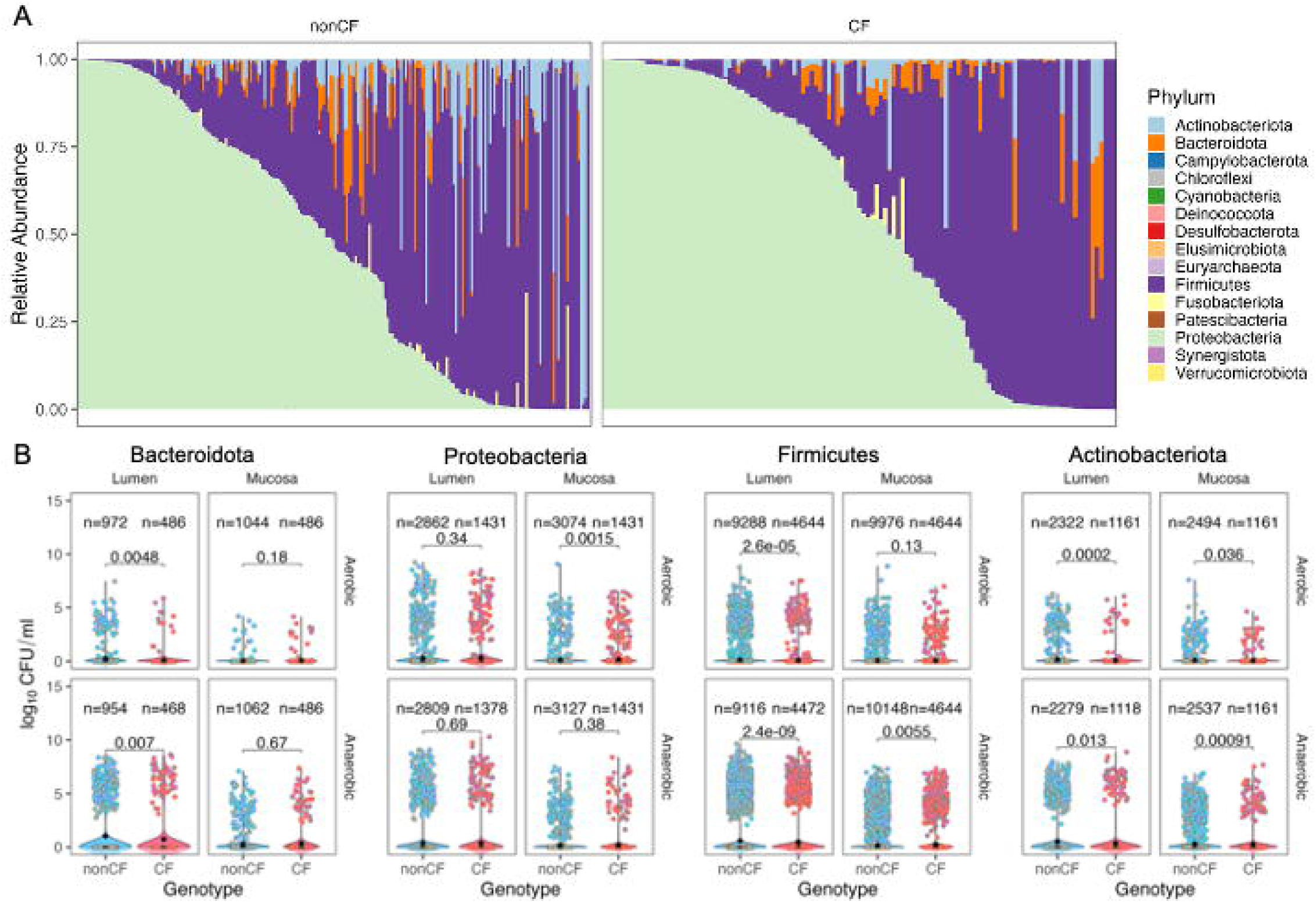
Bacterial composition and differences across major phyla. (A) Relative abundance of bacterial taxa in all samples, faceted by genotype (CF vs. non-CF) and colored by phylum. (B) Absolute abundance of the top four phyla, compared across microhabitat (luminal vs. mucosal) and oxygen status (top: aerobic; bottom: anaerobic). Black diamonds indicate group means. Colored dots are annotated by colonic location. Statistical comparisons were made using the Wilcoxon test for each category.

Because culturome sequencing and CFU counts were performed under the same culture conditions, we calculated (pseudo)absolute abundances by multiplying the relative abundance of each amplicon sequence variant (ASV) by the CFU/ml of the corresponding sample. Absolute abundance values of many phyla were significantly different (Fig. S5). This approach enabled direct quantification of bacterial loads at the taxonomic level. The effect size was calculated by subtracting the mean of group 1 from group 2 for each variable. Sample type influenced the largest number of phyla, as 8/10 phyla had a significantly higher abundance in luminal samples compared to mucosal. This observation aligns with the higher CFU load observed in the luminal samples. Another observation was a higher abundance of Bacteroidota in individuals with BMI >30, consistent with previous studies (25, 26). Desulfobacterota had the most impact based on sample type, with luminal samples having 0.71 log10 CFU/ml higher load than mucosal samples. This is consistent with literature reporting a greater abundance of sulfate-reducing bacteria in lumen/fecal contents under non-disease conditions, although some studies show mucosal enrichment in inflammatory settings (27, 28).

Comparison of absolute abundances between all CF and nonCF samples (i.e., genotype) revealed a significant decrease in Actinobacteria (effect size: -0.07 log10 CFU/ml), Bacteroidota (effect size: -0.06 log10 CFU/ml), and Firmicutes (effect size: -0.02 log10 CFU/ml; Fig. S5, S6) in pwCF. Proteobacteria abundance was elevated in pwCF but did not reach statistical significance (effect size: 0.04 log10 CFU/ml). To further assess how the major phyla respond to host and culture variables changes in CF, we focused on the four most abundant phyla: Bacteroidota, Proteobacteria, Firmicutes, and Actinobacteria (Fig. 4B). CF samples exhibited significantly reduced absolute abundance of Bacteroidota in luminal samples under both aerobic and anaerobic conditions, elevated Proteobacteria in aerobic mucosal samples, reduced Firmicutes in luminal samples under both oxygen conditions but increased Firmicutes in anaerobic mucosal samples, and consistently lower Actinobacteria across all sample types and conditions. These findings mirror stool-based reports of reduced Firmicutes, Bacteroidota, and Actinobacteria in CF, while revealing novel microniche-specific shifts that are only apparent in colonic subenvironments.

### Predictive Modeling of CF and Host Variables Using Cultured Communities

Given the significant differences in beta diversity across sample and patient variables, we evaluated the predictive potential of the culturome using random forest models. Models were trained with 10,000 trees to predict antibiotics use, BMI, colonic region, CFTR, genotype, GI symptoms, pathology, PPI use, probiotics, smoking, oxygen exposure during culture, and sample type.

All models achieved an out-of-bag (OOB) accuracy greater than 75%, except for the colonic region, which performed poorly (OOB accuracy: 22.3%; Table S6, Supplementary Data). For each model, the mean decrease in accuracy was calculated for each genus, and the top 15 predictive taxa were visualized (Fig. 5A). Genera including *Streptococcus*, *Enterococcus*, *Clostridium sensu stricto*, *Eubacterium*, and *Veillonella* consistently ranked among the top five predictive taxa across all models except for the colonic region.

**Figure 5.**
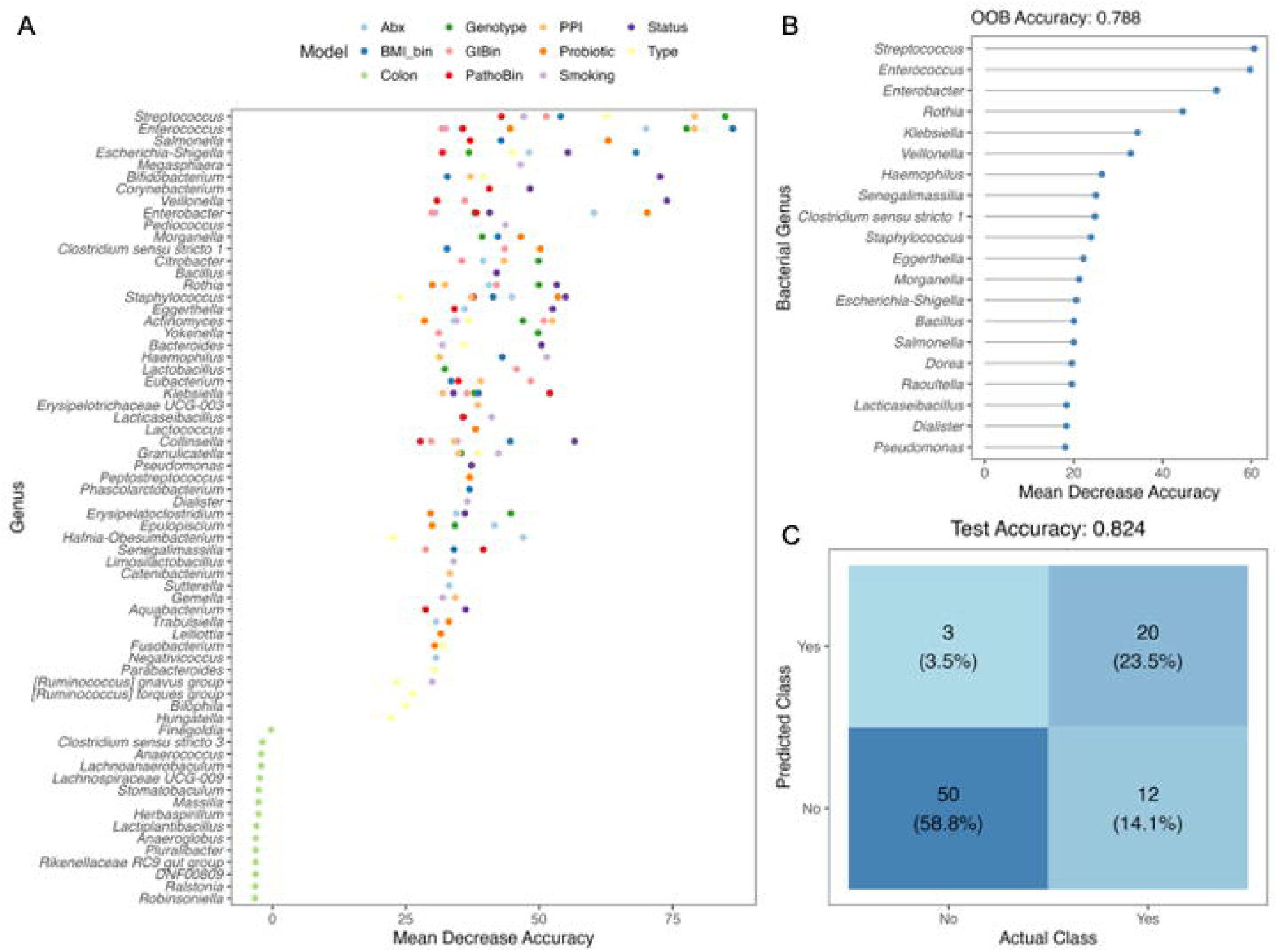
Multiple clinical outcomes can be predicted using culturome profiles. (A) Mean decrease accuracy of the top 15 genera for random forest models of the major clinical outcomes. (B) Mean decrease accuracy of the top 20 genera for training datasets of the random forest model for the presence of polyps. (C) Confusion matrix of the test dataset for the prediction of polyps using the polyp model from B. Models- Abx: Antibiotic use, BMI_bin: BMI categories; Colon: Colonic location, Genotype: CF or nonCF, GIBin: binary output of GI symptoms, PathoBin: binary output of diagnosed pathology, PPI: use of proton pump inhibitors, Probiotic: use of probiotics, Smoking: status of smoking, Status: Oxygen status of samples, Type: microhabitat of the sample.

To further validate the predictive performance, we randomly partitioned the dataset, using 75% for model training using the polyp information and 25% as an independent test set. The trained model for polyps achieved an OOB accuracy of 78.8%, with *Streptococcus*, *Enterococcus*, *Enterobacter*, *Rothia*, and *Klebsiella* identified as the most important predictors shown here as the mean decrease in accuracy (Fig. 5B). *Streptococcus* and *Rothia* abundance were decreased in CF, whereas *Enterococcus*, *Enterobacter,* and *Klebsiella* were higher in abundance (data not shown). When applied to the test dataset, the model predicted polyp status with 82.4% accuracy (Fig. 5C), demonstrating the potential utility of culturome-derived features as a biomarker.

### Strain-Level Genomic Variation Across the Colon

To further investigate strain-level variation, colonoscopy samples were streaked on a variety of selective media, and isolated colonies were subjected to whole-genome sequencing. From these strains, we selected twenty-one *Bacteroides fragilis* strains from GIEHZ100 (a heterozygous individual) and fifteen *Escherichia coli* strains from GEHV103 (a nonCF individual) for downstream genomic analyses. These isolates represent a subset of the 109 gut bacterial genomes sequenced from this cohort (22), enabling independent pangenome analysis of patient-specific strains.

Independent pangenomes were constructed for the patient-specific isolates to assess genomic variation across colonic regions and sample types. The pangenome of *B. fragilis* (Fig. S7A) and *E. coli* (Fig. S7B) revealed minimal differences among strains, regardless of their colonic origin (ascending, transverse, or descending) or sample type (mucosal vs. luminal). Core genome content was largely conserved, with the accessory genome showing only minor variability, suggesting that within an individual, these gut-associated strains are highly homogeneous at the genomic level.

## Discussion

This pilot culturome analysis reveals significant alterations in the CF colonic microbiome that extend beyond traditional fecal microbiome studies to include functionally relevant, culturable bacterial communities. Our findings demonstrate distinct spatial and phenotypic patterns of microbial dysbiosis in CF, with implications for understanding disease pathogenesis and therapeutic interventions.

The most striking finding of our study was the selective enrichment of viable bacterial load in CF mucosal samples, with no corresponding differences in luminal communities. This spatial specificity suggests that CF-associated microbial alterations are most pronounced at the host- microbe interface, where CFTR dysfunction directly impacts epithelial cell function (4–6). The mucosal-specific nature of these changes aligns with the known pathophysiology of CF, such as viscous mucous production, frequent exposure to antibiotics, altered bile acids, increased inflammation, increased fats, and atypical colonization patterns, resulting in an evolving dysbiosis of the gastrointestinal microsystems (29–31). Our findings of altered mucosal bacterial composition in CF adults align with the evidence that gut dysbiosis in CF begins early in life and has lasting consequences (11–13). This early establishment of dysbiosis may predispose CF individuals to the persistent mucosal bacterial alterations we observed in our adult cohort. The increased bacterial adherence to CF mucosal surfaces likely reflects both the altered physicochemical properties of CF mucus and impaired host antimicrobial defenses.

Microbiome studies have shown dysbiosis in children and adults with CF, either using samples from stool or small intestine (3, 10, 11, 13, 32). These previous studies report consistently decreased Bacteroidetes and increased Proteobacteria in children, with variable observations in adults. The taxonomic pattern observed in children directly parallels our culturome findings in adults, suggesting that the fundamental dysbiosis signature we detected represents a persistent feature of CF gut pathophysiology that emerges in infancy and continues throughout life. Our observations extend these known alterations to the colon, indicating that CFTR-mediated effects on mucosal bacterial communities are conserved throughout the gastrointestinal tract. These alterations can also be evident in our lack of changes observed across the colon. The functional implications of this dysbiosis extend beyond simple bacterial enumeration, as the affected taxa perform functions important for nutrient harvest and growth hormone signaling, potentially explaining the sustained impact on CF health outcomes (33).

The introduction of CFTR modulator therapies has revolutionized CF treatment, with drugs like elexacaftor/tezacaftor/ivacaftor providing unprecedented improvements in lung function and quality of life (34). However, the effects of these therapies on the gut microbiome remain incompletely understood. Extended use of CFTR modulators has been shown to shift the bacterial composition toward a “healthier” gut microbiome, but does not restore the endogenous communities (35, 36). Understanding how CFTR modulators affect gut microbial communities is crucial for several reasons: first, it may help explain some of the gastrointestinal side effects associated with these therapies; second, it could identify opportunities for combination therapies that optimize both CFTR function and microbial homeostasis; and third, it may reveal biomarkers for therapeutic response or guide personalized treatment approaches. Our observation that pwCF on modulators exhibited both higher bacterial loads and increased alpha diversity represents a paradoxical finding that requires careful interpretation. The increased bacterial diversity associated with modulator therapy may reflect restoration of a more hospitable gut environment, allowing previously suppressed bacterial taxa to re-establish. However, the concurrent increase in bacterial loads suggests that modulator therapy may also enhance overall bacterial growth conditions.

Our machine learning studies revealed the ability to predict polyp status with 82.4% accuracy using culturome data. In this model, we did not find any previously reported genera with high importance, such as *Bacteroides*, *Bifidobacterium*, *Fusobacterium, Faecalibacterium*, and *Blautia,* typically associated with dysbiosis in pwCF (37). This finding might be due to either a small sample size or culturing bias. We also detected higher bacterial load in individuals with diagnosed pathology using CFUs. With more validation, this predictive capacity, along with higher viable loads, could potentially enable earlier intervention or more targeted surveillance strategies for pwCF at risk for gastrointestinal complications. Our finding is also clinically significant because these culturable organisms could serve as readily assessable biomarkers for disease monitoring and therapeutic responses.

The relevance of oral microbes to gut microbiome composition in CF has important implications for interpreting our results. The oral cavity represents an additional component of this microbial network, and emerging evidence indicates that the oral cavity acts as a bacterial reservoir and might contribute to the transmission of bacteria to the GI tract in diseases (38). Several of the predictive genera we identified in our machine learning models, such as *Streptococcus*, *Megasphaera, Veillonella,* and *Corynebacterium,* are commonly found in the oral cavity and could potentially represent oral-gut translocation. The presence of these oral-derived taxa in the gut culturome could contribute to the mucosal-specific bacterial elevation we observed, as these organisms may preferentially adhere to mucosal surfaces after translocation from the oral cavity.

Our study also generated a strain collection from multiple individuals and colonic locations. These strains belong mostly to gut bacterial genera such as *Bacteroides*, *Clostridium*, *Escherichia*, *Enterococcu*s*, Lactococcus, Lactobacillus, and Thomasclavelia,* along with oral microbes like *Streptococcus*. Patient-specific single-species pangenome analysis showed the presence of clonal isolates consistent with previous observation (39). These findings indicate that, despite environmental differences along the colon or between mucosal and luminal niches, the dominant strains of *B. fragilis* and *E. coli* within a given individual maintain a remarkably stable genomic composition.

Several limitations of our study should be acknowledged. First, our culturome approach, while providing insights into functionally relevant bacterial communities, represents only a fraction of the total gut microbiome diversity. Second, the colonoscopy preparations might also alter the gut composition. Third, the relatively small sample size, particularly for pwCF, may limit the generalizability of some findings; however, the observation of robust and significant differences in the CF and nonCF cohorts, which support previous findings, speaks to the magnitude of the differences. Future studies should incorporate longitudinal sampling to assess the stability and dynamics of culturome changes over time, particularly in response to modulator therapy.

Additionally, integration of culturome data with metabolomic and immunological analyses could provide deeper insights into the functional consequences of microbial alterations in CF.

## Acknowledgments

This work was supported by the Cystic Fibrosis Foundation grant OTOOLE22G0 and the NIH grant R01-ES033988. We acknowledge samples, facilities, and expertise offered by Dartmouth’s Translational Research Core and Gastrointestinal Biology Core (NIH/P30-DK117469), specifically Deborah A. Hogan. We would like to thank Daniel J. Gutzmann, Ana Victoria de Sousa Bezerra, and Davon L. Gokey for their help with DNA extractions.

## Data Availability

Amplicon sequencing reads have been deposited at NCBI under BioProject accession number PRJNA1299459. Genome assemblies and raw sequencing reads have been deposited at NCBI under BioProject accession number PRJNA833080.

## Notes

### Competing Interest Statement

The authors have declared no competing interest.

