## Supplemental Figures S1-7 for "The Regional Landscape of the Human Colon Culturome in Health and Cystic Fibrosis"

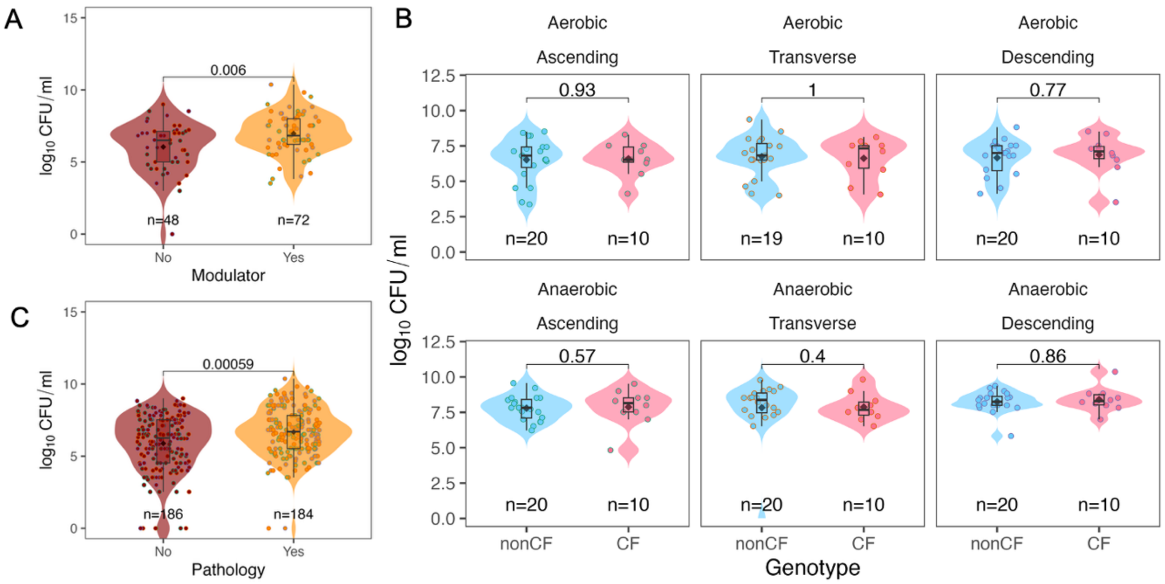


**Figure S1. No differences in CFUs in luminal samples.** Viable counts ($log_{10}$ CFUs) comparing (A) use of modulators, and (B) genotype for luminal samples only, stratified by colonic region (ascending, transverse, descending) and oxygen status (top: aerobic; bottom: anaerobic), and (C) diagnosed pathology. In all plots, black diamonds indicate group means. Colored dots are annotated by colonic location. Statistical comparisons were made using the Wilcoxon test for each category.

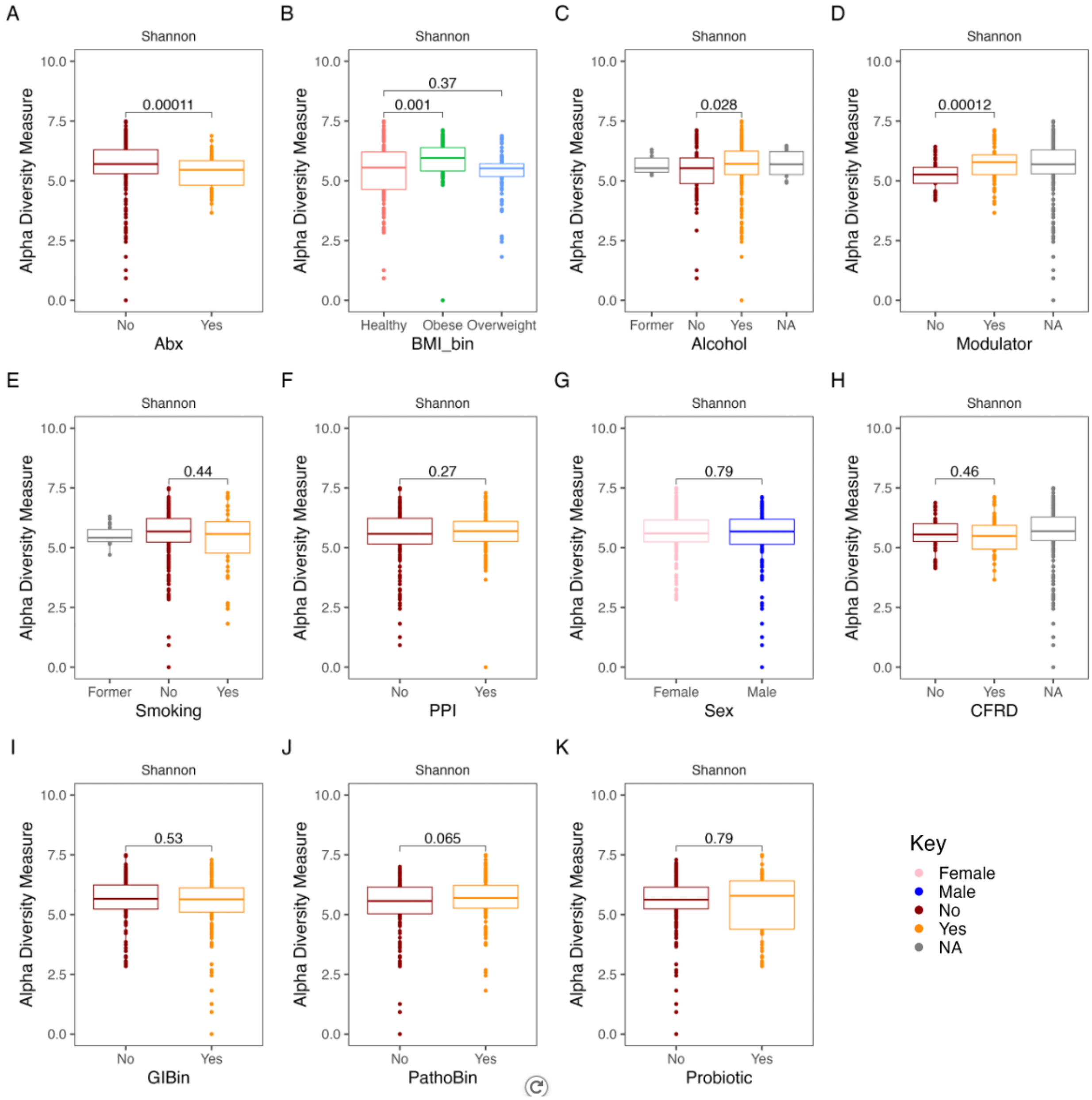


**Figure S2. Some lifestyle variables change bacterial diversity.** Alpha diversity of all samples, calculated using the Shannon Diversity Index (SDI) comparing (A) antibiotic use, (B) BMI categories, (C) alcohol use, (D) modulator use in case of CF, (E) status of smoking (F) intake of proton pump inhibitors, (G) sex of the individual, (H) status of CFRD for CF individuals, and binary outputs of (I) GI symptoms, (J) diagnosed pathology, and (K) probiotic use. For the variables CFRD and Modulator, 'NA' indicates non-CF individuals; in all other cases, 'NA' indicates information not available. Statistical comparisons were performed using the Wilcoxon test. 'NA' indicates information not available.

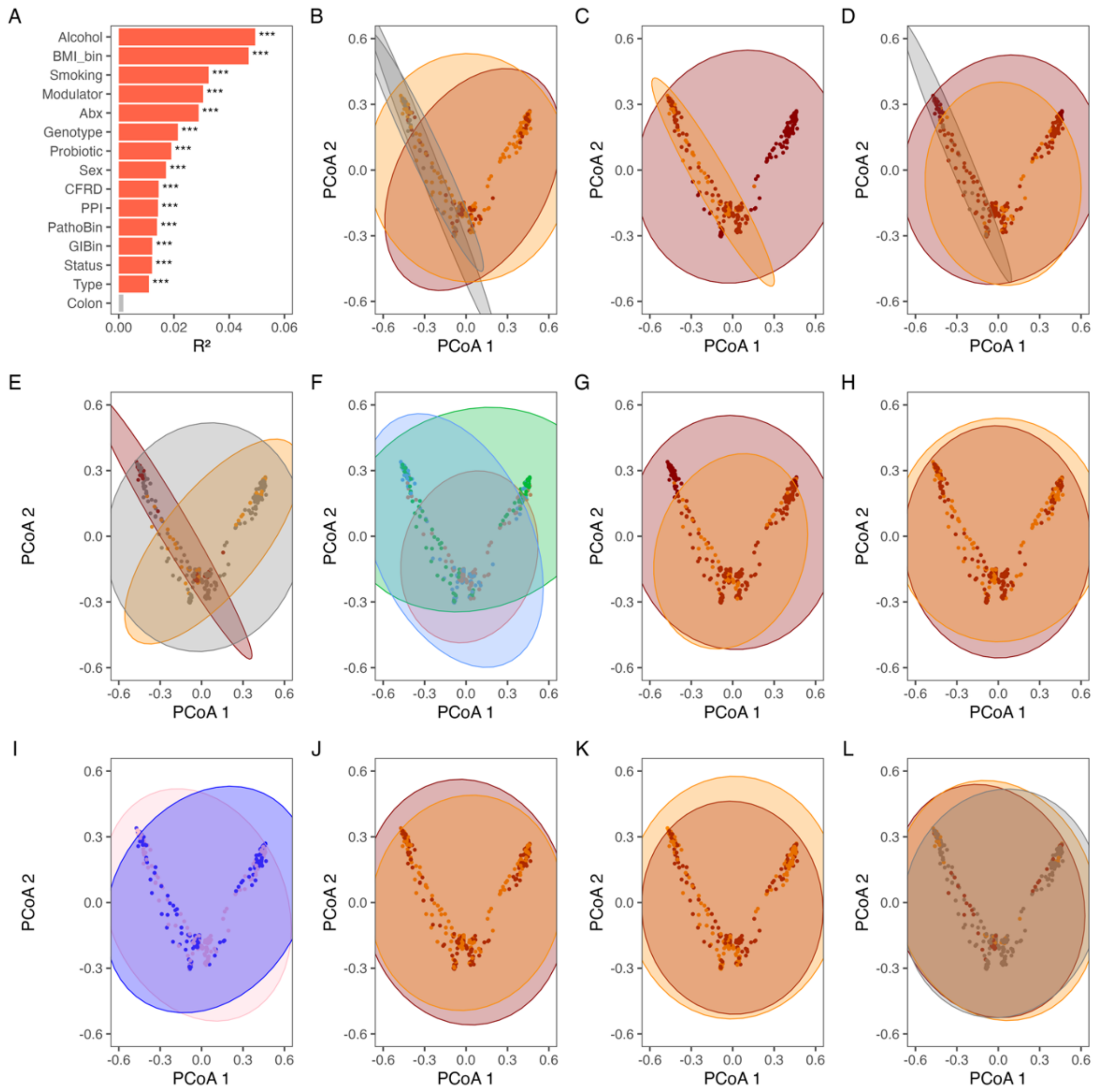


**Figure S3. Some lifestyle variables change bacterial diversity.** (A) PERMANOVA $R^2$ and p-values for each of the tested variables. Beta diversity, based on Bray-Curtis dissimilarity and visualized using Principal Coordinates Analysis (PCoA) comparing (B) alcohol use, (C) antibiotic use, (D) status of smoking, (E) modulator use in case of CF, (F) BMI categories, (G) probiotic use, (H) intake of proton pump inhibitor, (I) sex of the individual, binary outputs of (J) diagnosed pathology, and (K) GI symptoms as mentioned in Table S1, and (L) status of CFRD for CF individuals. Color key is the same as Figure S2. For the variables CFRD and Modulator, 'NA' indicates non-CF individuals; in all other cases, 'NA' indicates information not available.

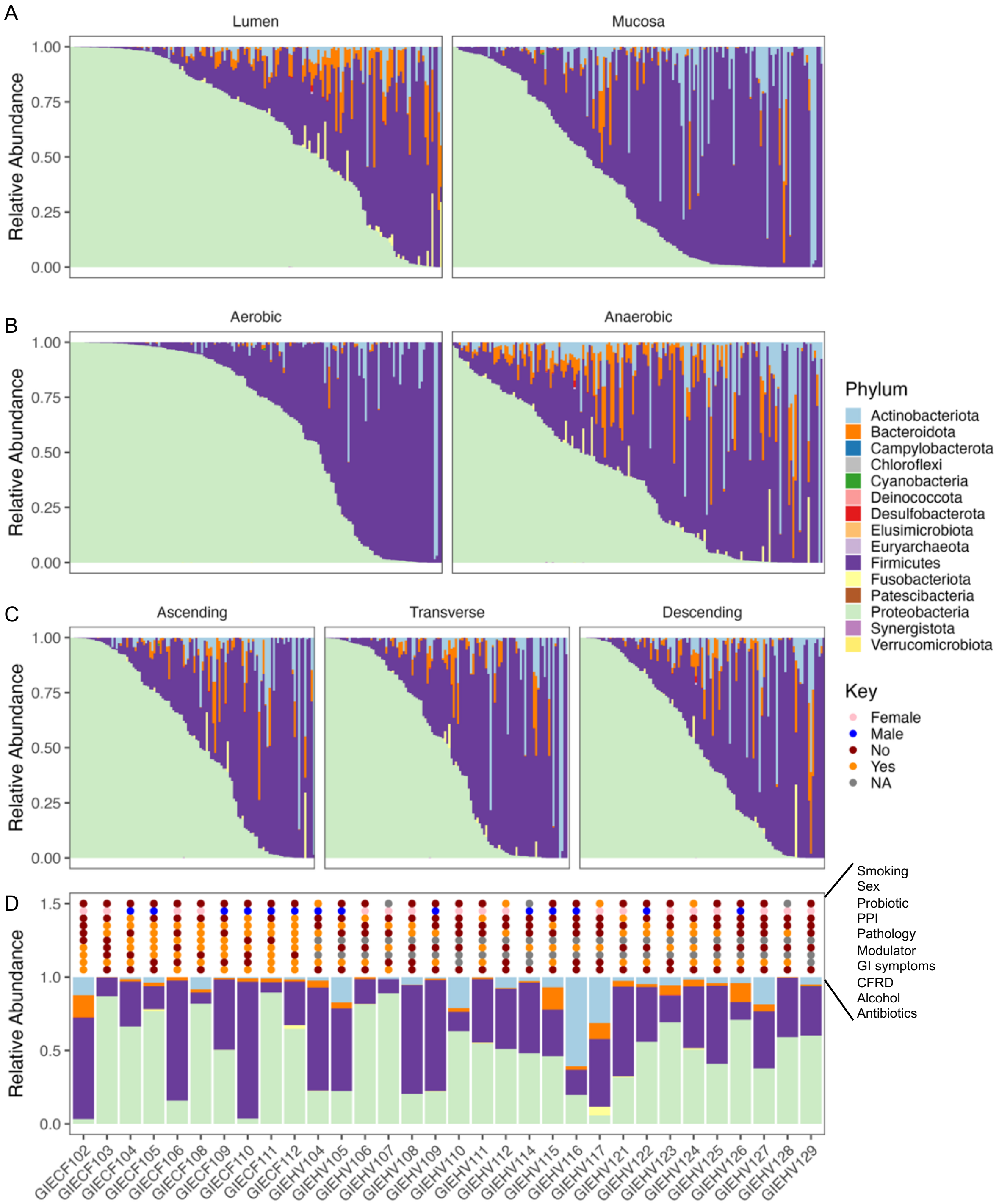


**Figure S4. Bacterial composition in different conditions**. Relative abundance of bacterial taxa in all samples colored by phylum and faceted by (A) microhabitat (lumen and mucosa), (B) oxygen status (aerobic and anaerobic), and (C) colonic location (ascending, transverse, and descending). (D) The bacterial community profiles per patient, showing the mean of all the conditions with host factors colored according to the key on top.

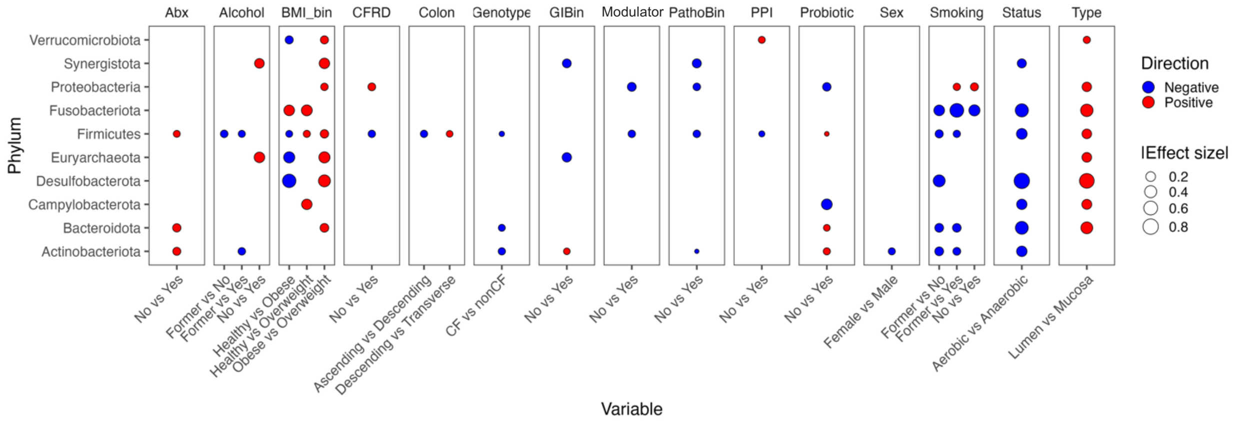


**Figure S5. Enrichment of phyla in different conditions.** Pairwise comparison (group 1 vs group 2) of absolute abundances for each variable using the Wilcoxon test, showing only the statistically significant ($p<0.05$) phyla. The size of the dot shows the absolute effect size, and the color shows the direction of enrichment. An increase in group 1 abundance is shown in red, whereas an increase in group 2 is shown in blue. Antibiotic use, Alcohol: use of alcohol, BMI_bin: BMI categories; CFRD: CFRD status for CF individuals, Colon: Colonic location, Genotype: CF or nonCF, GIBin: binary output of GI symptoms, Modulator: use of modulators for CF individuals, PathoBin: binary output of diagnosed pathology, PPI: use of proton pump inhibitors, Probiotic: use of probiotics, Sex: biological sex of the individual, Smoking: status of smoking, Status: Oxygen status of samples, Type: microhabitat of the sample.

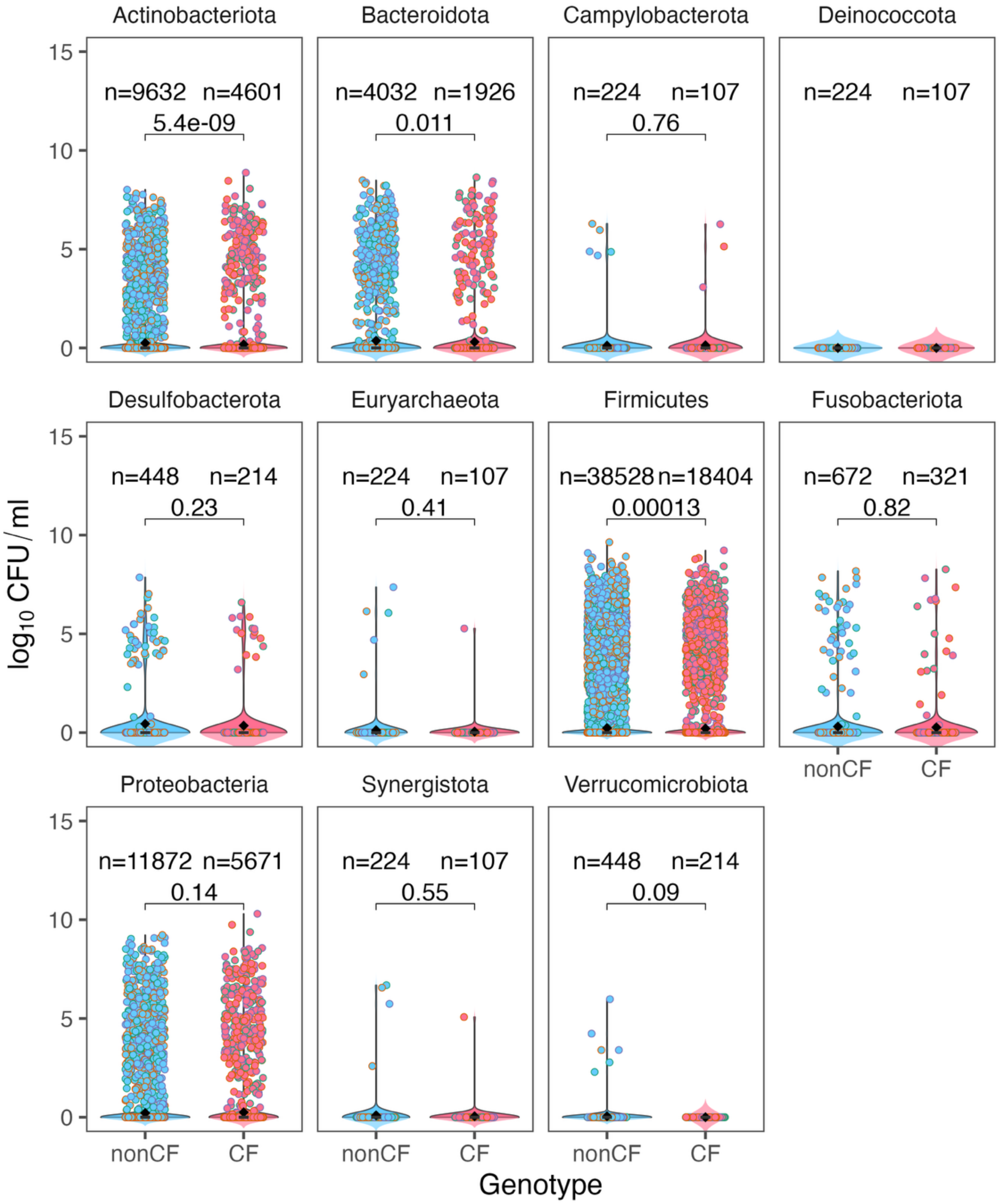


**Figure S6. Absolute abundance changes in CF individuals across phyla.** Viable counts ($log_{10}$ CFUs) comparing CF and non-CF genotypes, faceted by phyla. In all plots, black diamonds indicate group means. Colored dots are annotated by colonic location. Statistical comparisons were made using the Wilcoxon test for each category.

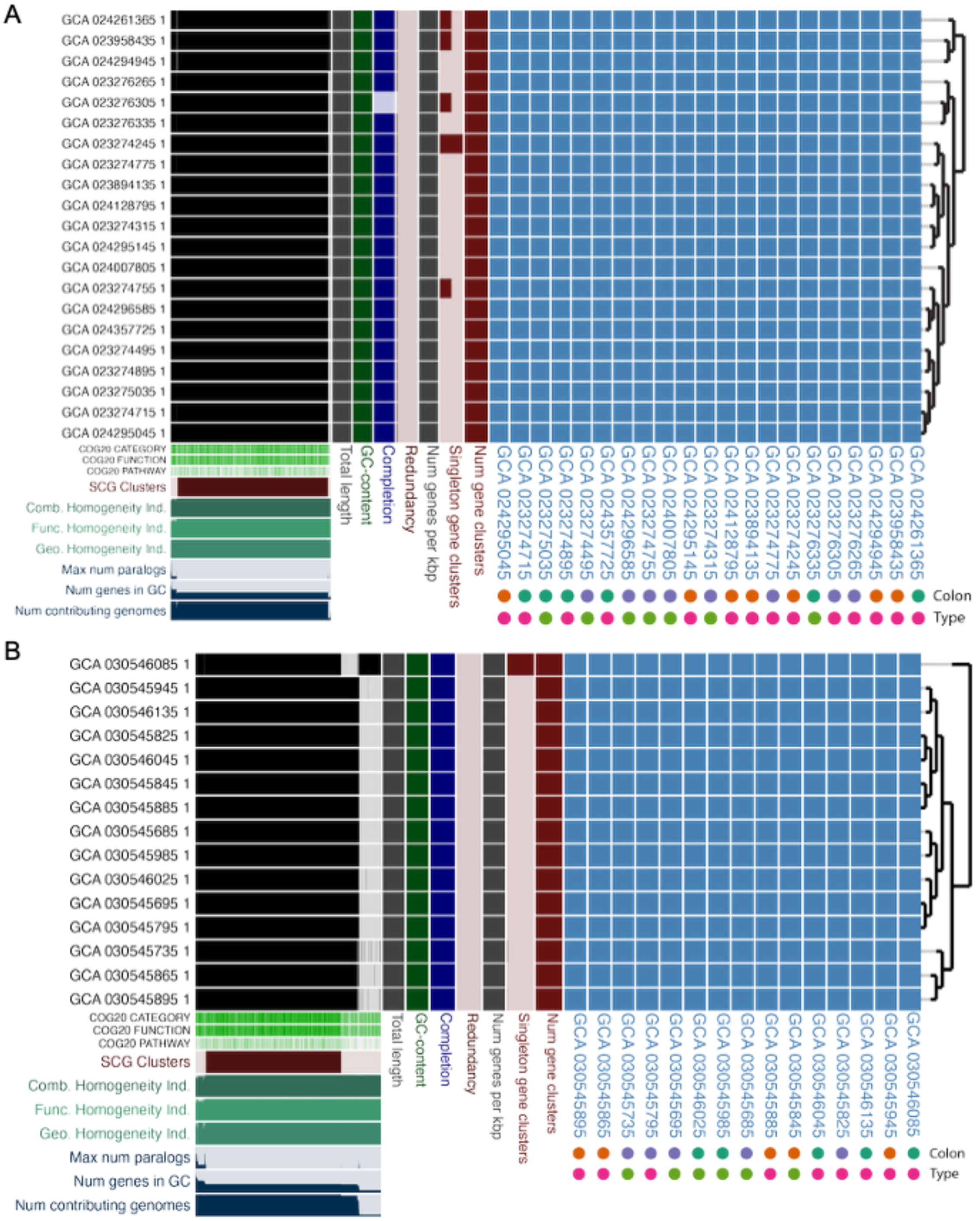


**Figure S7. Pangenomes of *Bacteroides fragilis* and *Escherichia coli* isolates from a single patient.** Pangenomes of (A) *B. fragilis* and (B) *E. coli* recovered from the ascending (green), transverse (orange), and descending (violet) colon, and from luminal (pink) and mucosal (green) samples. Gene cluster presence-absence matrices are shown on the left with associated metadata below and to the right. Average nucleotide identity (blue) is shown on the right.
